# Recognition memory: familiarity signals increase selectively in the lateral entorhinal cortex following hippocampal lesions

**DOI:** 10.1101/2020.06.16.131763

**Authors:** Liv Mahnke, Erika Atucha, Takashi Kitsukawa, Magdalena M. Sauvage

**Affiliations:** Leibniz-Institute for Neurobiology, Functional Architecture of Memory Dept., 39118, Magdeburg, Germany; Otto von Guericke University, Medical Faculty, Functional Neuroplasticity Dept., 39120, Magdeburg Germany; Otto von Guericke University, Center for Behavioral Brain Sciences, 39106, Magdeburg Germany; KOKORO-biology group, Osaka University, 565-0871, Osaka, Japan

## Abstract

The sense of familiarity for events is crucial for successful recognition memory. However, the neural substrate and mechanisms supporting familiarity remain unclear. Some human and animal studies suggest that the lateral entorhinal (LEC) and the perirhinal (PER) cortices might be essential for familiarity judgments while others attribute this function to the hippocampus (HIP) and it is unclear whether LEC, PER and HIP interact within this frame. Here, we especially investigate if LEC and PER’s contribution to familiarity depends on hippocampal integrity. Using a human to rat translational memory task and high resolution IEG imaging, we report that hippocampal lesions selectively enhance activity in LEC during familiarity judgments. These findings suggest that different mechanisms support familiarity in LEC and PER and, that HIP might exert a tonic inhibition on LEC during recognition memory that might be released when HIP is compromised, possibly constituting a compensatory mechanism in aging and amnesic patients.

## Introduction

The medial temporal lobe (MTL) includes the hippocampus as well as brain areas surrounding the hippocampus that are crucial for memory function: the parahippocampal areas^1, 2^. The lateral (LEC) and medial entorhinal (MEC) cortices are part of the parahippocampal areas as well as the peri- and postrhinal cortices (PER and POR, respectively). Decades of studies have investigated the role of the hippocampus (HIP) in memory function in humans and animals while empirical data on the parahippocampal regions, especially on LEC and MEC, have started accumulating only recently with the discovery of the grid cells in animals^3, 4^ and the functional dissociation of LEC and MEC in humans^5, 6^. Some early memory studies have however predicted that LEC might support the recognition of familiar items (i.e. single objects, odors etc…) mainly based on its strong anatomical ties with PER^7, 8^, while others have attributed this function to HIP in addition to its well-accepted role in the recollection of episodic events^9, 10, 11^. Plethora of studies have reported a specific involvement of PER in the familiarity process, especially during spontaneous object recognition memory in rodents and word recognition memory in humans^12, 13, 14, 15, 16, 17, 18, 7, 8^. In contrast, clear empirical evidence for a specific involvement of LEC in familiarity judgements is lacking in humans, essentially because human studies typically investigate BOLD signal in the anterior parahippocampal gyrus reflecting the activation of both PER and LEC.^8^ Likewise, evidence for a role of the LEC in familiarity in animals is very scarce with, to date, only one study reporting a clear contribution of this area using a response deadline design yielding familiarity judgments. ^17^ This is a clear shortcoming as conflicting reports suggest that LEC and PER might not constitute a single functional entity.^19, 20, 21^ Thus, the extent to which familiarity truly relies on LEC function remains unclear.

Importantly, LEC shares major direct bidirectional projections with HIP that has been suggested by some to support familiarity judgments in addition to recollection^9, 10, 11^, while PER and HIP are much less anatomically connected^22, 23, 24^. In addition, the extent to which HIP might affect PER function has been extensively investigated especially during object recognition memory in animals^25, 26^. However, no study has yet investigated whether hippocampal function might influence LEC’s contribution to familiarity judgments. Hence, the extent to which the familiarity signal in LEC might be tied to hippocampal activity and if this relationship is comparable to that between PER and HIP is not known.

To address these questions, we studied activity patterns in LEC and PER in rats with hippocampal lesions, known to rely on familiarity to solve a translational human to rat delayed non-matching to odor recognition memory task (Fig. 1A)^27^, and compare these patterns to those observed in rats with intact hippocampal function known to rely on both familiarity and recollection to solve the same task^27^. To do so, we imaged brain activity during memory retrieval using a high-resolution imaging technique (i.e. to the cellular level) based on the detection of the immediate-early gene (IEG) *Arc* RNA. This technique is commonly used to map MTL activity and allows for the assessment of the percentage of cells recruited during cognitive tasks^28, 29, 30, 31, 32, 33, 34, 35^. These comparisons revealed a robust and comparable recruitment of LEC and PER during familiarity judgments as well as a selective increase of activation of LEC following HIP lesions within this frame (i.e. not of PER), suggesting that distinct mechanisms might support familiarity in these areas.

**Figure 1A:**
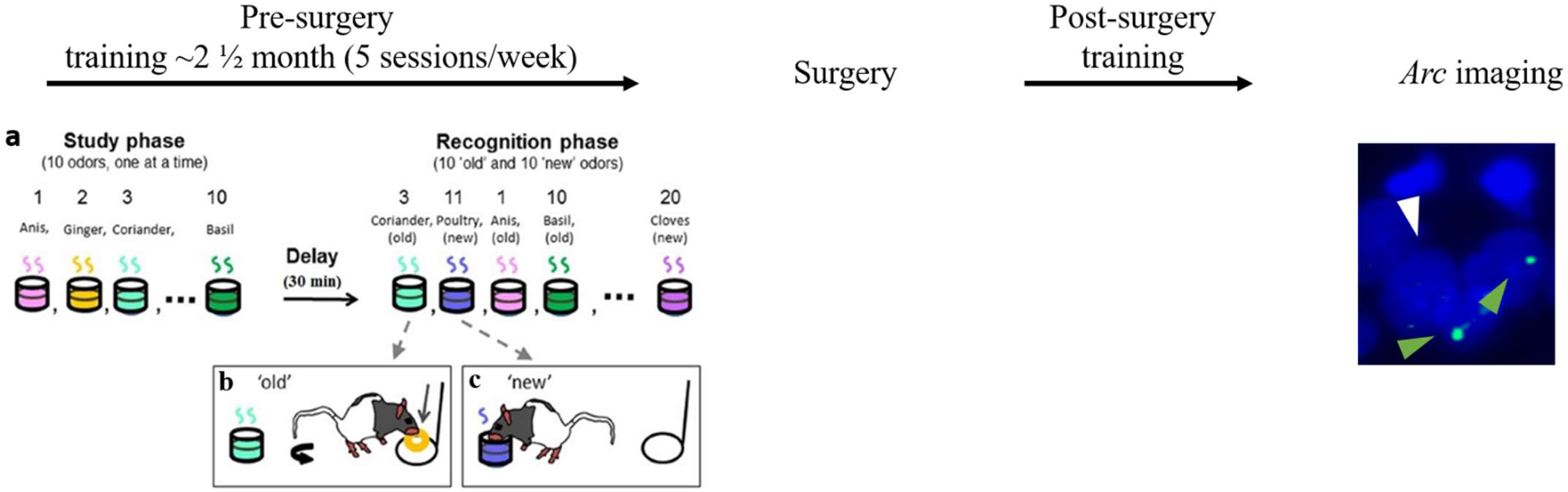
Overview of the experimental design. *Left*: Pre-surgery training. **(a)** Ten odors are presented to the animal during the study phase (one at a time). After a 30 min delay, the memory for the studied odors is tested by presenting the same odors intermixed with 10 new odors to the animals (also one at a time). **b**, **c:** Delayed nonmatching-to-sample rule. If the odor belonged to the study list (old odor), the rat was expected to refrain from digging and turn around to get a food reward at the back of the cage **(b)**. If the odor did not belong to the study list (new odor), the rat could dig in the stimulus cup to retrieve a buried reward **(c)**. *Right:* After reaching the criterion, HIP and sham-surgeries were performed. After recovery, post-surgery training lasted until rats reached a plateau performance on 3 consecutive days subsequently to which rats were sacrificed and their brain processed for *Arc* imaging (green arrowheads: examples of *Arc* positive cells, white arrowhead: *Arc* negative cell, nuclei are counterstained with DAPI). [Adapted from Nakamura and Colleagues^38^]

## Method and Materials

### Subjects and stimuli

Adults male Long Evans rats (350-430g) were maintained under reverse light/dark cycle (7:00 A.M. light off/7:00 P.M. light on), food deprived up to 85% of their body weight and received water *ad libitum*. The animals were handled a week before the experiment and tested in their home cage. The stimulus odors were common household scents (anise, rosemary, fennel, etc.) mixed with playground sand and this scented sand (one odor per cup) was held in glass cups (HHIndustries). A pool of 40 scents was available and 20 pseudo randomly chosen odors were used each day. All procedures were approved by the animal care committee of the State of Saxony-Anhalt (42502-2-1555 LIN) and performed in compliance with the guidelines of the European Community.

### Behavioral paradigm

Behavioral training followed the training protocol previously described in Sauvage et al.^36, 37^. In brief, to study recognition memory, we used the innate ability of rats to dig and to discriminate between odors. Each training session contained a study phase, a delay, and a recognition phase (Fig. 1A). On each daily training session rats were presented with a unique study list of 10 odors, i.e. the list changed for each session. Following a 30-min retention delay, memory was tested using a list of odors that included the 10 odor stimuli (old odors) used during the study phase intermixed with 10 additional odors that were part of the pool of forty odors but were not presented during the study phase (new odors; Fig. 1Aa). During the recognition phase of the task, animals were tested for their ability to distinguish between the 10 odors that were presented during the study phase (old odors) and the 10 additional odors (new odors). Animals were first trained to dig in the stimulus cup with unscented sand to retrieve one ¼ of piece of Honey loops (Kellogg’s) and were subsequently trained on a delay non-matching-to-sample (DNMS) rule. During the recognition phase, when rats were presented with an odor that was part of the study list (an old odor), rats were required to refrain from digging, turn around, and go to the back of the cage to receive a food reward: a correct response for an old odor (Fig. 1Ab; an incorrect response would be digging in the stimulus cup). Conversely, when the odor was not part of the study list (a new odor), animals could retrieve a buried reward by digging in the test cup: a correct response for a new odor (Fig. 1Ac; an incorrect response would be going to the back of the cage to receive the reward). To ensure that the task could not be solved by smelling the reward buried in the sand, all cups were baited but the reward was not accessible to the animal for the old odors. In addition, no spatial information useful to solve the task was available to rats, given that testing cups for new and old odors were presented at the exact same location. Reward locations differed for the new and old odors (front and back of the cage, respectively), but were only experienced by the animals once a decision had been made (e.g., when the trial was over), hence could not contribute to behavioral performance. Training lasted ~3½ months including surgery recovery time and post-surgery training. Pre-surgery training consisted of several steps during which the number of studied odors increased from one to 10, the delay increased from one to 30 minutes and the number of odors during the recognition phase increased from two to 20 (half old, half new). The post-surgery training mirrored the final stage of the pre-surgery training (10 odors study list, a 30 min delay and a 20 odors testing list). Animals transitioned between successive training stages when performance reached a minimum of 80% correct for three consecutive days. After reaching the final training stage (10 study odors, 30 min delay, and 20 test odors) and performing at least 80% correct for three consecutive days, the animals were split in two groups of equivalent memory performance. Subsequently, animals underwent surgery and received either a selective lesion to the hippocampus (the HIP lesion group) or a sham-surgery (the HIP intact group). After 2 weeks of recovery, rats were trained until a plateau performance was reached over 3 consecutive days and sacrificed immediately after completion of the last recognition phase, which lasted ~8 min. Throughout the training, each HIP lesion rat was paired with an intact HIP rat of comparable pre-surgery performance and both animals were sacrificed on the same day. Also, to ensure that *Arc* premRNA expression was low at baseline, brain activity was also imaged in an additional home-caged control group that was exposed to the same testing room as the performing animals (i.e. placed in the same room throughout the training), but to which no memory demands was applied.

### Surgery

Rats were anaesthetized by using 0,5% pentobarbital (diluted in 1,2-Propanediol) and placed into a stereotactic frame (Kopf Instruments, Tujunga, CA, USA). A heating pad was used to control body temperature, and eyes were coated with a moisturizing balm. Additional pentobarbital was applied to maintain anesthesia, when necessary. The lesions were made by injecting N-Methyl-D-aspartic acid (10 μg/μL NMDA in 0.9% saline; Sigma, Germany) into the dorsoventral and mediolateral hippocampus at 8 sites bilaterally at a flow rate of 0.15 μL/min (see (Table 1 for coordinates). The injection needle was left in place for an additional 2.5 minutes following the injection to facilitate diffusion and then slowly withdrawn. The sham surgery rats (HIP intact group) underwent the same surgical procedure (craniotomy and placement of the needle) but no NMDA was injected. All animals were allowed to recover for two weeks before the post-surgery training took place. Upon lesion assessment using Nissl‘s staining (Fig. 3A), one HIP lesion rat was placed in the HIP intact group as hippocampal damage was minimal (ca. 17%), no change between pre-and post-surgery memory performance was observed and adding this animal in the HIP intact did not alter differences found in memory performance nor in IEG patterns of activity in LEC and LER between HIP intact and HIP lesioned groups. Thus, the final group size for the HIP lesion group was n=6 against n=7 for the HIP intact group.

**Table 1.** Coordinates relative to Bregma (in mm) and corresponding volumes (in μl) for NMDA injections in the hippocampus

| Anterior-posterior<br>(AP) | Medio-lateral<br>(ML) | Dorso-ventral<br>(DV) | Volume per<br>site (μL) |
| --- | --- | --- | --- |
| -3.6 | $\pm 1.0$ | -3.6 | 0.2 |
| -3.6 | $\pm 2.0$ | -3.6 | 0.2 |
| -4.6 | $\pm 2.0$ | -4.0 | 0.2 |
| -4.6 | $\pm 3.5$ | -4.0 | 0.2 |
| -5.5 | $\pm 3.0$ | -4.1 | 0.15 |
| -5.5 | $\pm 5.2$ | -6.3 | 0.15 |
| -5.5 | $\pm 5.2$ | -4.0 | 0.15 |
| -6.3 | $\pm 4.4$ | -4.4 | 0.15 |
| -6.3 | $\pm 5.1$ | -6.5 | 0.15 |
| -6.3 | $\pm 5.1$ | -5.5 | 0.15 |

### Brain collection

Animals were deeply anesthetized with isoflurane and decapitated. Brains were immediately collected, frozen in isopentane cooled in dry ice, and subsequently stored at −80°C. Brains were then coronally sectioned on a cryostat (8μm sections; Leica CM 3050S, Leica Microsystems), collected on polylysine-coated slides, and stored at −80°C.

### Fluorescent *in situ* hybridization histochemistry

The *Arc* DNA template was designed to amplify a fragment containing two intron sequences from bases 1934–2722 of the rat *Arc* gene (NCBI Reference Seq: NC_005106.2). DIG-labeled *Arc* RNA probes were synthesized with a mixture of digoxigenin-labeled UTP (DIG RNA Labeling Mix, Roche Diagnostics) and purified using Probe quant G-50 Micro columns (GE Healthcare). Fluorescent *in-situ* hybridization histochemistry was performed as previously described in the study of Nakamura et al.^38^ with modifications. In brief, slides were fixed with 4% buffered paraformaldehyde and rinsed several times with 0.1 M PBS. After washing, the slides were treated with an acetic anhydride/triethanolamine/hydrochloric acid mix (0.25% acetic anhydride in 0.1 M triethanolamine/HCl), rinsed with 0.1 M PBS and briefly soaked with a prehybridization buffer. The prehybridization buffer contained 50% formamide, 5×SSC, 2.5×Denhardt’s solution, 250 μg/ml yeast tRNA and 500 μg/ml denatured salmon sperm DNA. For the hybridization step a 0.05 ng/μl digoxigenin-labeled *Arc* RNA probe was applied to each slide and after adding a coverslip, the slides were incubated in a humidified environment at 65°C for 17 h. After the hybridization, sections were first rinsed at 65°C in 5×SSC and then in 0.2×SSC for 1 h. Subsequently the slides were rinsed in 0.2×SSC at room temperature before the sections were incubated with 1% bovine serum albumin (BSA) in TBST buffer (0.1 M Tris-HCl pH 7.4, 0.15 M NaCl, 0.05% Tween 20) for 15min. The incubation with the anti-digoxigenin-POD took place afterwards (1/2000 dilution in BSA/TBST, Roche Diagnostics) at room temperature for 3 h. Sections were then rinsed with TBST and the signal amplified using the Tyramide Signal Amplification (TSA) Cy5 System. DAPI (4’,6’-diamidino-2-phenylindole, 1/100.000) was used to counterstain nuclei. Slides were then coverslipped and stored at +4°C. Additional slides without *Arc* pre-mRNA probe were used as a negative control. In these slides, no *Arc* labelling was detected.

### Image acquisition

To image *Arc*, one slide per animal was processed. Slides contained 4 nonconsecutive brain sections and images from nonadjacent sections distant approximately 200 microns were acquired. The number of activated neurons was evaluated on approximately 55 neurons per area on 3 nonadjacent sections (i.e., on a total of approximately 165 neurons per area covering approximately 400 microns). Images were captured with a Keyence Fluorescence microscope (BZ-X710; Japan). Images were taken with a 40× objective (Nikon) for PER and LEC and a 20×objective for CA1 and CA3 in z-stacks of 0.7-μm–thick images containing 12 to 16 images (see example Fig. 2B). The exposure time, light intensity, contrast and gain settings were kept constant between image-stacks. As first described in the seminal work of Guzowski and colleagues^39^, contrasts were set to optimize the appearance of intranuclear foci^40, 41, 42, 43^. LEC, PER, CA1 and CA3 images were captured at the anteroposterior (AP) levels: −5.1 to −5.3 mm defined from Bregma (Fig. 2A)^41^. Hippocampal ROIs (only in HIP intact rats) were defined based on HIP areas showing maximal *Arc* expression in the same task^17^.

### Counting of *Arc*-positive cells

To account for stereological considerations, neurons were counted on 8-μm–thick sections that contained one layer of cells, and only cells containing whole nuclei were included in the analysis^44^. The quantification of *Arc* expression was performed in the median 60% of the stack in our analysis because this method minimizes the likelihood of taking into consideration partial nuclei and decreases the occurrence of false negative. This method is comparable to an optical dissector technique that reduces sampling errors linked to the inclusion of partial cells into the counts and stereological concerns because variations in cell volumes no longer affect sampling frequencies^45^. Also, as performed in a standard manner in *Arc* imaging studies, counting was performed on cells (>5μm) thought to be pyramidal neurons or interneurons because small non-neuronal cells such as astrocytes or inhibitory neurons do not express *Arc* following behavioral stimulation^46^. The designation “intranuclear-foci–positive neurons” (*Arc*-positive neurons) was given when the DAPI-labeled nucleus of the presumptive neurons showed 1 or 2 characteristic intense intranuclear areas of fluorescence. DAPI-labeled nuclei that did not contain fluorescent intranuclear foci were counted as “negative” (*Arc*-negative neurons) (see Fig.2B for examples of *Arc*-positive and *Arc-*negative images)^39^. Cell counting was performed manually by experimenters blind to experimental conditions. Percentage of *Arc*-positive neurons was calculated as follows: *Arc*-positive neurons / (*Arc*-positive neurons + *Arc*-negative neurons) × 100.

### Statistics

Data are expressed as mean ± SEM. Unpaired t-tests (two-tailed) were used to compare memory performance between HIP lesion and HIP intact groups and paired t-tests to compare pre- and post-surgery performance. One-sample t-tests were used for comparison to chance level (50%). To compare *Arc* expression two-tailed unpaired t-tests were used for between-groups comparisons (lesion vs intact) and two-tailed paired t-tests for within group comparisons LEC and PER). Given the a priori hypothesis, a one-tailed paired t-test was used to compare *Arc* expression inCA1 and CA3 in the HIP intact group. For comparisons to zero one-sample t-tests were used. Finally, a One-way ANOVA was used to compare baseline *Arc* expression between MTL areas in home caged control rats. All statistical analyses were performed using the software IBM SPSS Statistics version 23.

## Results

### Memory performance and assessment of hippocampal lesions

Animals learned to discriminate ‘old’ from ‘new’ odors over 49 ± 1 training sessions. Once the criterion was reached (at least 80% correct responses for 3 consecutive days), two groups of comparable memory performance were formed: one hippocampus (HIP) lesioned group and one HIP intact group (performance pre-surgery: HIP lesion: 85.8% ± 1.5, HIP intact: 84.3% ± 1.3, t_(11)_ = 0.77, p = 0.45; see Fig. 1B). Subsequently, a hippocampal lesion was performed on the lesioned group and a sham surgery on the intact group (i.e. the hippocampus was not lesioned in the latter group). After two weeks of recovery, both groups were trained until the HIP intact group reached again the criterion, which took 9±1 training sessions. Memory performance of the HIP intact group following surgery was comparable to performance prior to surgery (HIP intact pre- vs post-surgery: 84.3% ± 1.3 vs 85% ± 2.9, t_(6)_ = −0.20, p = 0.85). Conversely, as reported in the study of Fortin and colleagues 27 a significant drop in performance was observable in the lesioned group (HIP lesion pre- vs post-surgery: 85.8% ± 1.5 vs 68.3% ± 3.6, t_(5)_ = 4.13, p = 0.009) which was also observable when compared to the HIP intact group post-surgery performance (HIP lesion: 68.3% ± 3.6; HIP intact: 85% ± 2.9; t_(11)_ = −3.67, p = 0.004). Importantly, performance in both groups was higher than chance level indicating that the HIP lesion group was still successfully retrieving memories (comparisons to chance level (50% correct): HIP lesion: t_(5)_ = 5.13, p = 0.004; HIP intact: t_(6)_ = 12.12, p < 0.001), albeit with lower accuracy than the HIP intact group. Lesion assessment revealed that in average 75% ± 9,9 of the hippocampus was damaged in the HIP lesion group (see Fig. 3A and B).

**Figure 1B:**
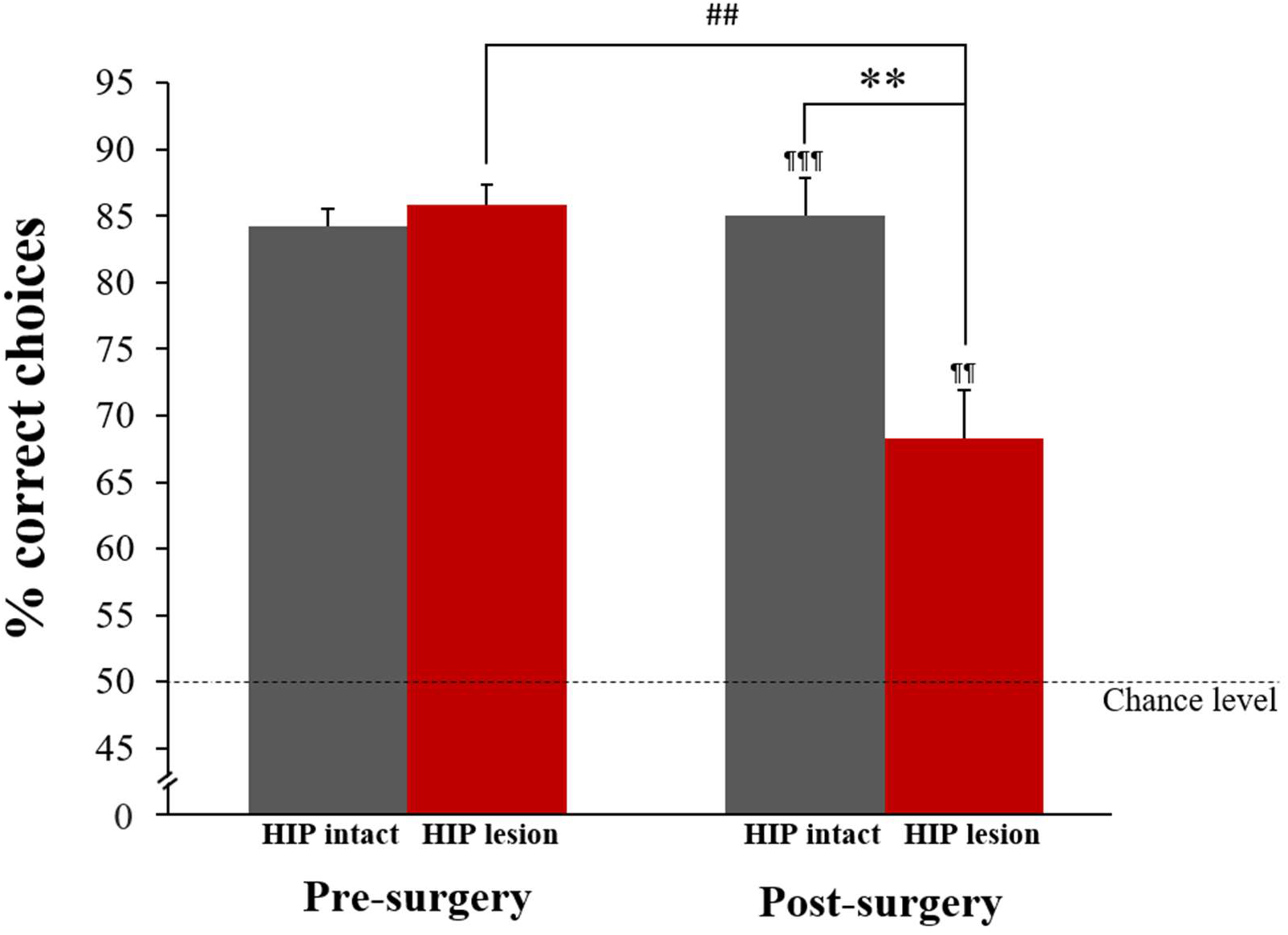
Pre-and post-surgery memory performance. Pre-surgery performance is comparable between HIP intact and lesioned groups. Likewise, pre and post- surgery performance did not differ for the intact HIP group. Lesion of HIP significantly altered memory performance as shown by a lower performance post- than pre- surgery in the HIP lesion group (^##^p < 0.01) and when compared to HIP intact performance post-surgery (**p < 0.01). Importantly, post-surgery, HIP lesion rats could still successfully perform the task, albeit with a lowest accuracy than HIP intact rats (comparison to chance level: ^¶¶^p < 0.01, ^¶¶¶^p < 0.001).

**Figure 2A:**
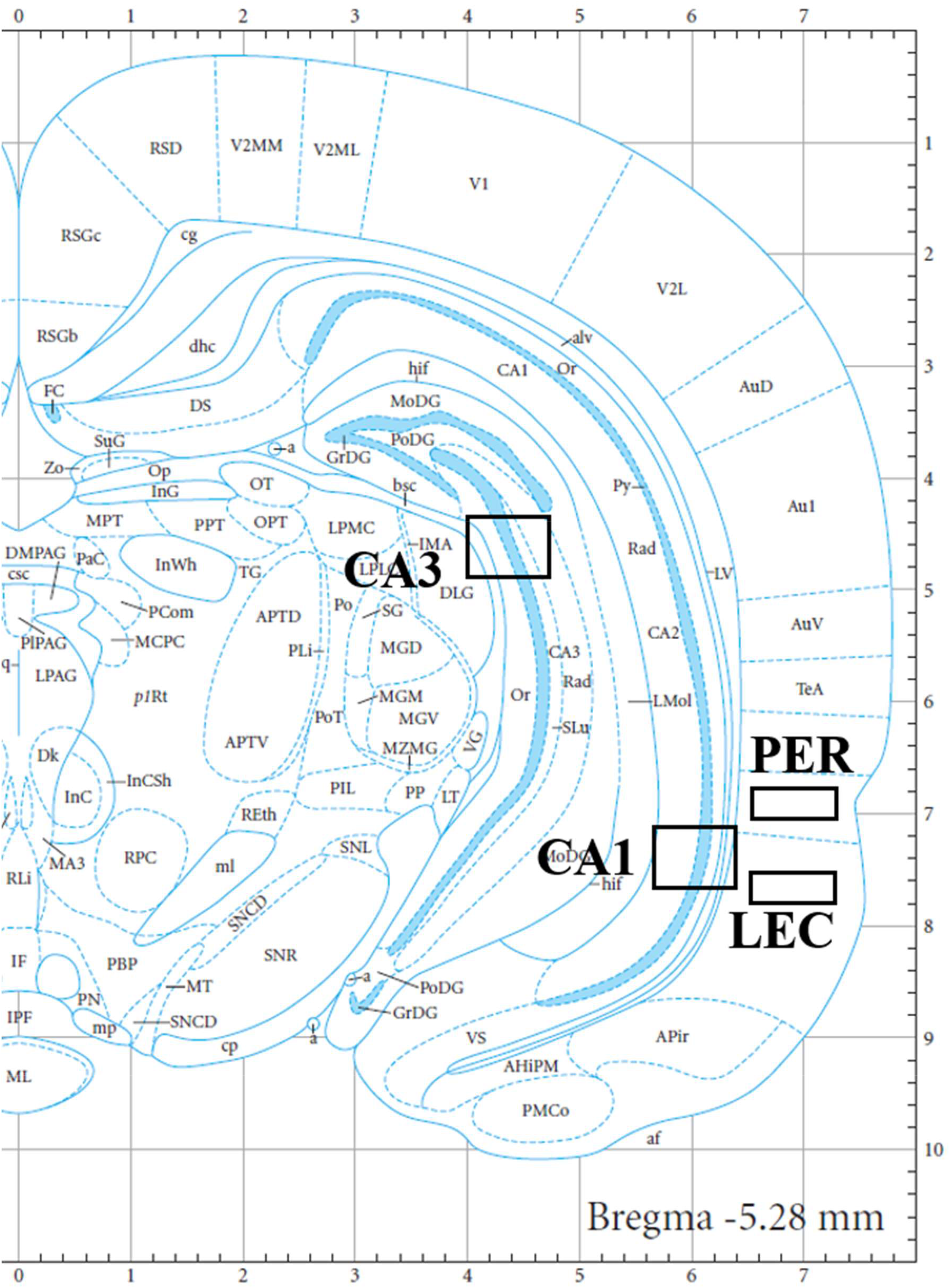
Location of the imaging frames for the regions of interest. CA1: cornu ammonis field 1; CA3: cornu ammonis field 3; PER: perirhinal cortex; LEC: lateral entorhinal cortex

**Figure 2B:**
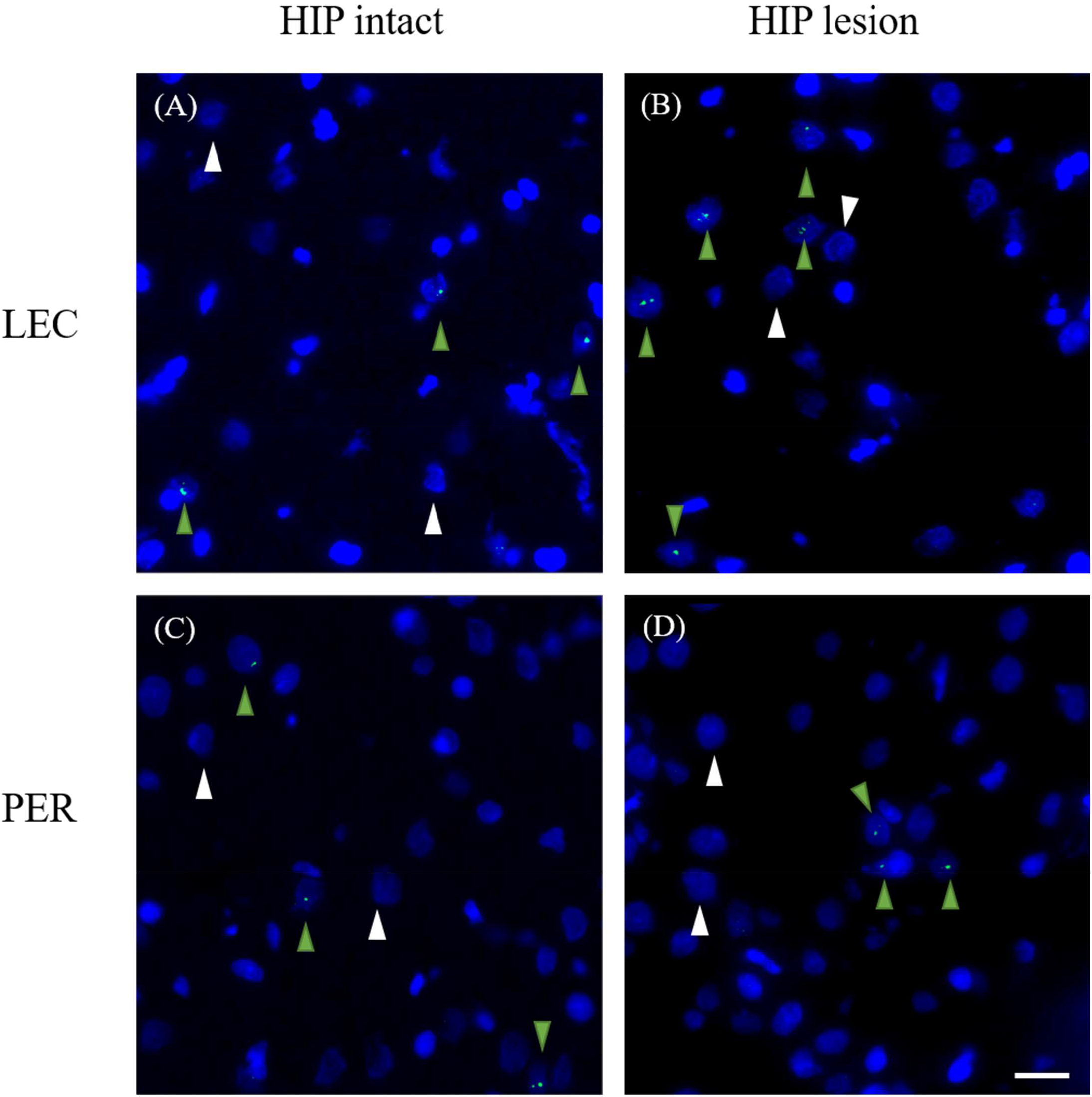
Representative images of *Arc* RNA expression in LEC and PER during memory retrieval in HIP intact and HIP lesioned groups. Lesioning HIP enhances the percentage of *Arc* positive cells in LEC (A vs B), while it has no significant effect in PER (C vs D). DAPI-stained nuclei are shown in blue. *Arc* intranuclear signal in green. Green arrowheads show *Arc* positive cells. White arrowheads show *Arc* negative cells. Scale bar 20 μm.

**Figure 3:**
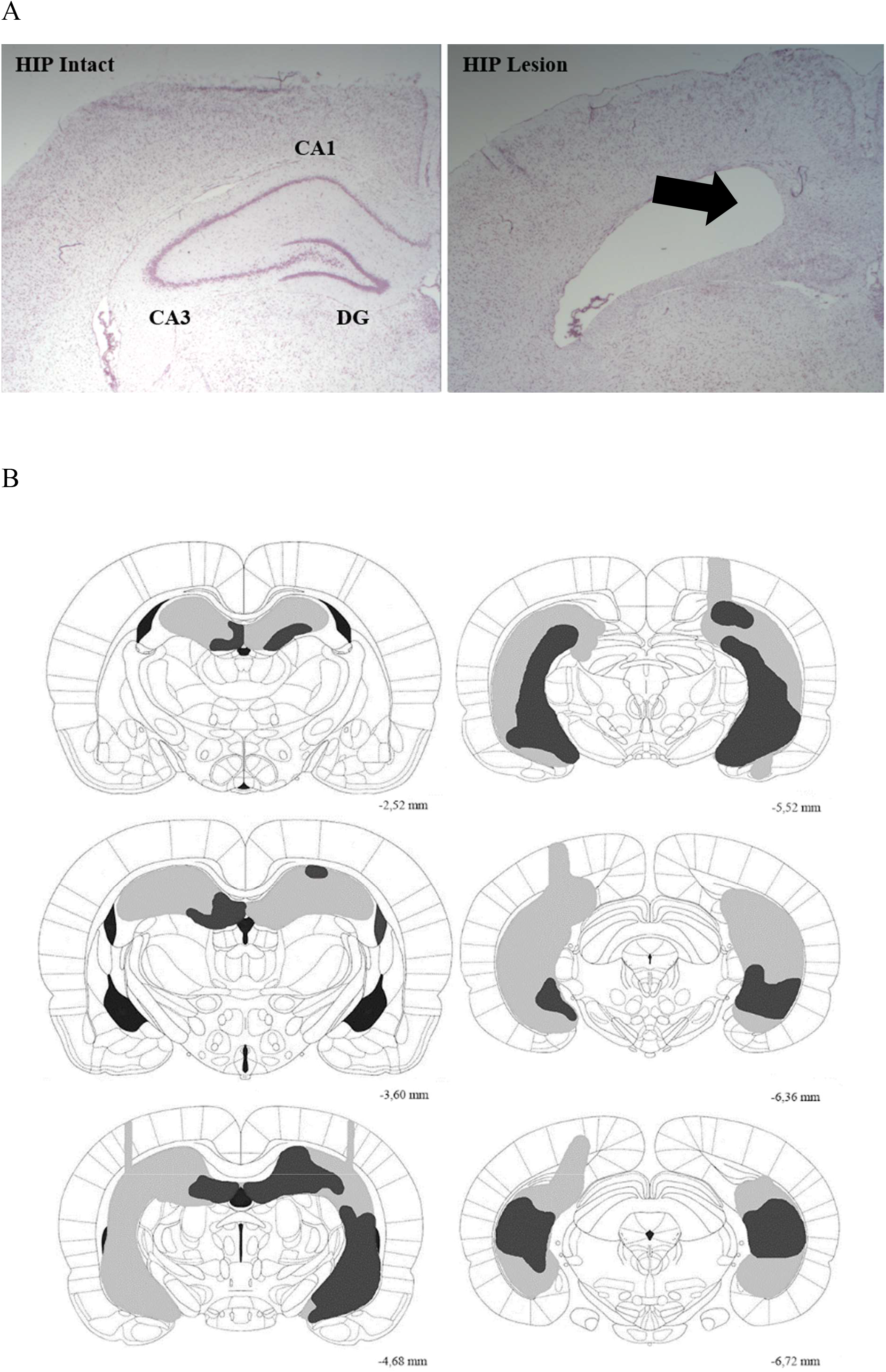
Extent of the HIP lesion. **A)** Photomicrographs of Nissl stained coronal sections of HIP intact (left) and HIP lesion (right) rats. **B)** Representation of the HIP lesion. Light grey: largest lesions. Dark grey: smallest lesions. Approximately 76% of HIP was lesioned in average.

### Hippocampal lesion affects familiarity signals in LEC, but not in PER

LEC and PER were engaged during the retrieval phase of the DNMS task in both the HIP intact and HIP lesion groups (comparisons to 0: HIP intact: LEC t_(6)_ = 8.87, p < 0.001; PER t_(6)_ = 9.15, p < 0.001; HIP lesion: LEC t_(5)_ = 9.93, p < 0.001; PER t_(5)_ = 8.06, p < 0.001; Fig. 4). In the HIP intact group, LEC and PER were recruited to a comparable level (LEC vs PER: t_(6)_ = 1.28, p = 0.25). In addition, further statistical comparisons showed for the first time that impaired hippocampal function led to an increase in LEC activity (HIP lesion vs intact groups: t_(11)_ = −2.53, p = 0.028; Fig. 4), revealing that LEC activity is sensitive to HIP lesion and that LEC is more recruited in rats relying primarily on familiarity (the HIP lesion group) than in rats relying on both familiarity and recollection (the HIP intact group). In sharp contrast, activity levels in PER remained comparable independently of whether hippocampal function was compromised or not (HIP lesion vs intact groups: PER: t_(11)_ = −1.15, p = 0.27; Fig. 4), indicating that PER activity is independent of hippocampal function within this framework. This differential effect of hippocampal lesion on LEC and PER activity was further supported by a direct comparison between LEC and PER showing a higher engagement of LEC than PER following hippocampal lesion during retrieval of odor memories (HIP lesion LEC vs PER: t_(5)_ = 3.67, p = 0.014), despite a comparable level of engagement in rats with intact hippocampus (HIP intact LEC vs PER: t_(6)_ = 1.28, p = 0.25). Thus, these results suggest an inverse functional relationship between HIP and LEC during familiarity judgments and the absence of strong ties between PER and HIP function within this frame.

**Figure 4:**
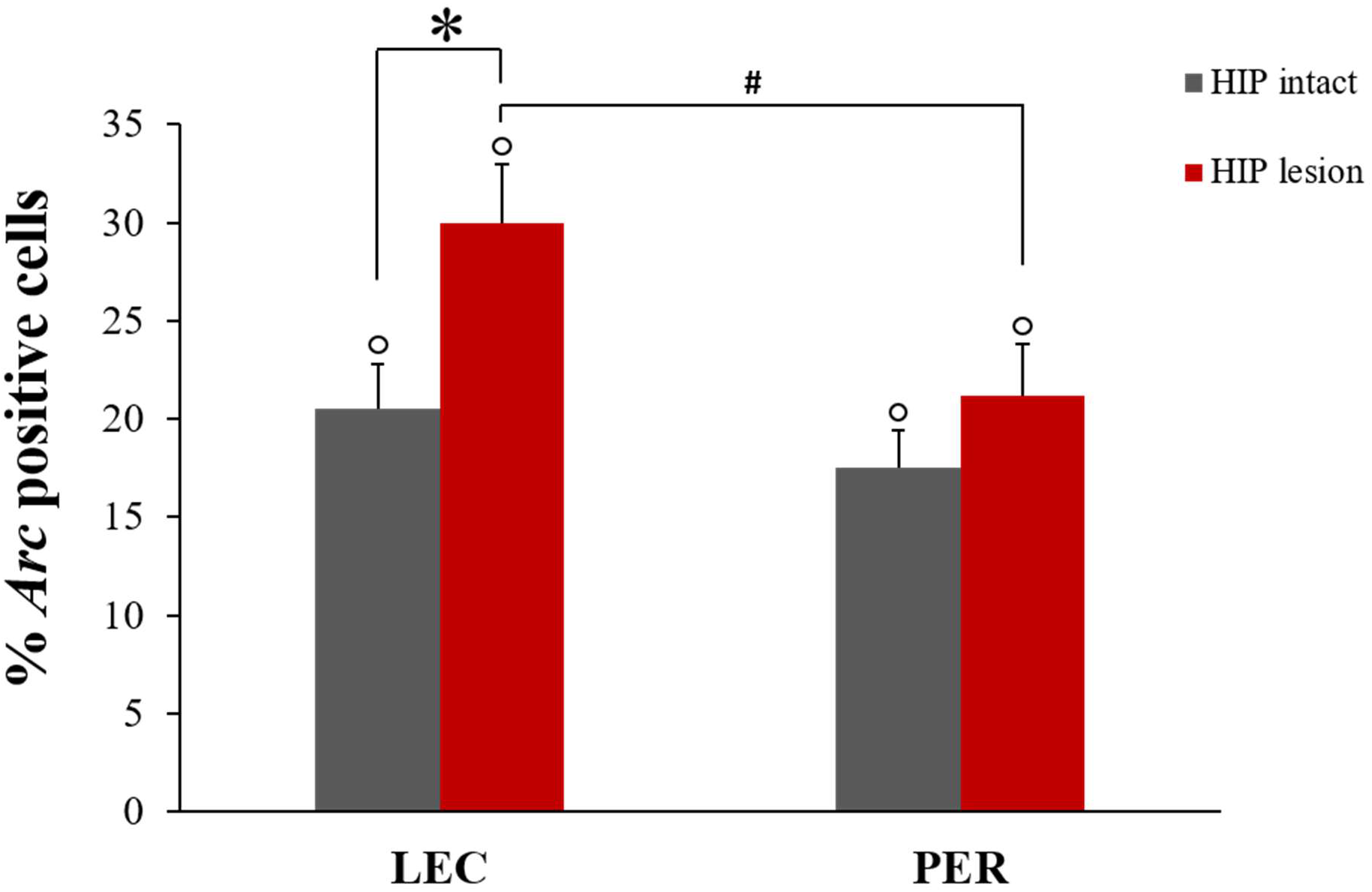
Percentage of *Arc* positive cells in LEC and PER in HIP intact and lesioned rats performing the memory task (bars represent means ± SEM). LEC and PER are recruited during the task in both groups (comparisons to 0: °p < 0.001). Importantly, *Arc* RNA expression is increased only in LEC and not in PER following HIP lesion (* p < 0.05), suggesting a distinct functional relationship between LEC and HIP, and PER and HIP. In addition, LEC is more recruited than PER during the task in rats with compromised hippocampal function (^#^ p < 0.05) while they are recruited to a similar extent in rats with intact hippocampus, suggesting a potential compensatory mechanism.

### Besides LEC and PER, HIP is also engaged during the task in rats with intact hippocampus

In addition to relying on familiarity, rats with intact functional hippocampus were shown to also rely on recollection (supported by the hippocampus) to solve the task used in the present study27. Thus, we controlled that the hippocampus was engaged during the task in rats with intact functional hippocampus (understandingly, such an assessment could not be performed in the HIP lesion group as HIP has been lesioned). High resolution imaging of CA1 and CA3 revealed a strong recruitment of these areas at retrieval, with a higher recruitment of CA3 than CA1, indicating that the hippocampus was indeed engaged in addition to LEC and PER in intact HIP rats (comparison to 0: CA3: t_(3)_ = 15.45, p = 0.001, CA1: 0: t_(3)_ = 6.68, p = 0.007; CA3 vs CA1: t_(3)_ = 2.76, p = 0.035; Fig. 5; see previous paragraph for LEC and PER statistical comparisons;).

**Figure 5:**
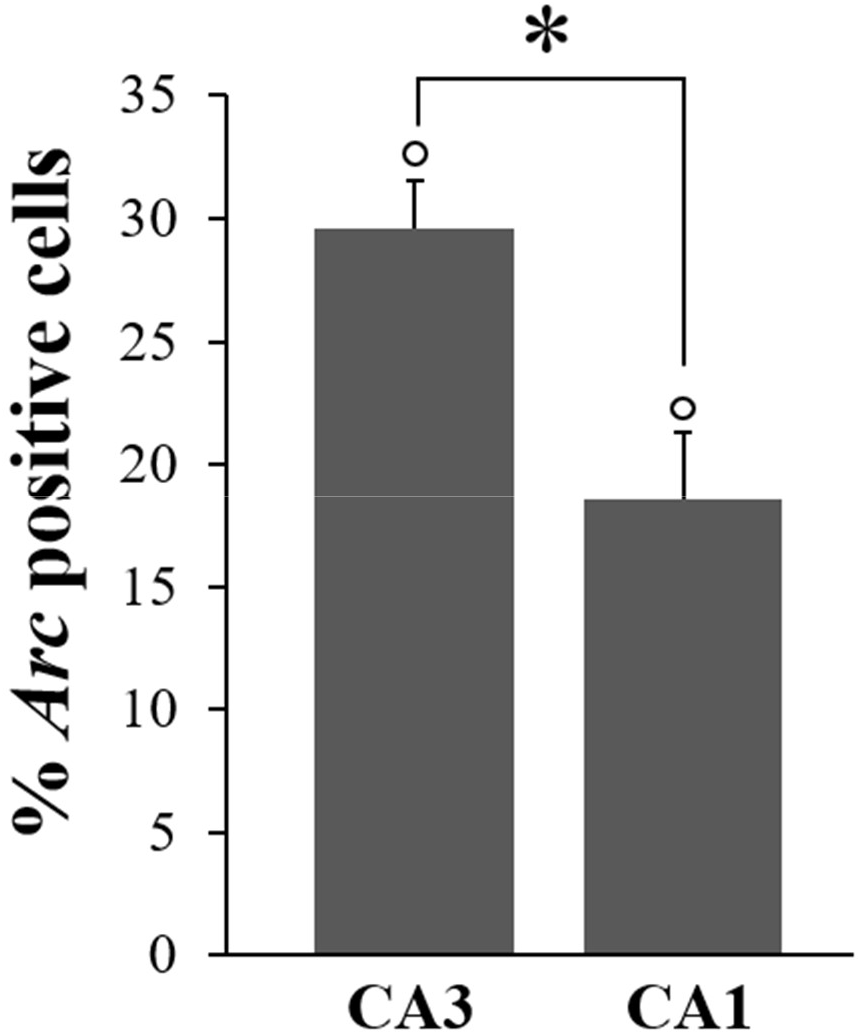
Percentage of *Arc* positive cells in CA3 and CA1 hippocampal subfields in HIP intact rats performing the memory task (means ± SEM). In addition to LEC and PER, CA3 and CA1 are highly recruited during memory retrieval in rats with intact hippocampus that rely on recollection and familiarity to solve this task (comparisons to 0: °p < 0.01; CA3 more than CA1: *p: < 0.05).

### LEC, PER, CA1 and CA3 are only mildly engaged in rats that do not perform the task

As expected baseline *Arc* RNA expression in LEC, PER, CA1 and CA3 of home-caged controls was low (LEC: 8.27% ± 2.5; PER: 6.54% ± 2.27, CA1: 5.37% ± 0.6, CA3: 12.63% ± 2.74; comparisons to 0: LEC t_(5)_ = 3.31, p = 0.021; PER t_(5)_ = 2.88, p = 0.035; CA1: t_(3)_ = 8.92, p = 0.003; CA3: t_(3)_ = 4.61, p = 0.019: Fig. 6) and comparable across areas (F _(3,16)_ = 1.55, p = 0.24; Fig. 6). These rats did not perform the task but were exposed to the testing room according to the same experimental scheme than rats performing the task. Also, as expected, *Arc* RNA expression in animals performing the task was overwhelmingly higher than in home-caged controls (HIP intact vs controls: LEC t_(11)_ = 3.59, p = 0.004; PER t_(11)_ = 3.72, p = 0.003; CA1: t_(6)_ = 4.64, p = 0.004; CA3: t_(6)_ = 5.08, p = 0.002; HIP lesion vs controls: LEC t_(10)_ = 5.54, p < 0.001; PER t_(10)_ = 4.21, p = 0.002), indicating that differences in *Arc* expression are likely to capture the cognitive demands of the task^39^.

**Figure 6:**
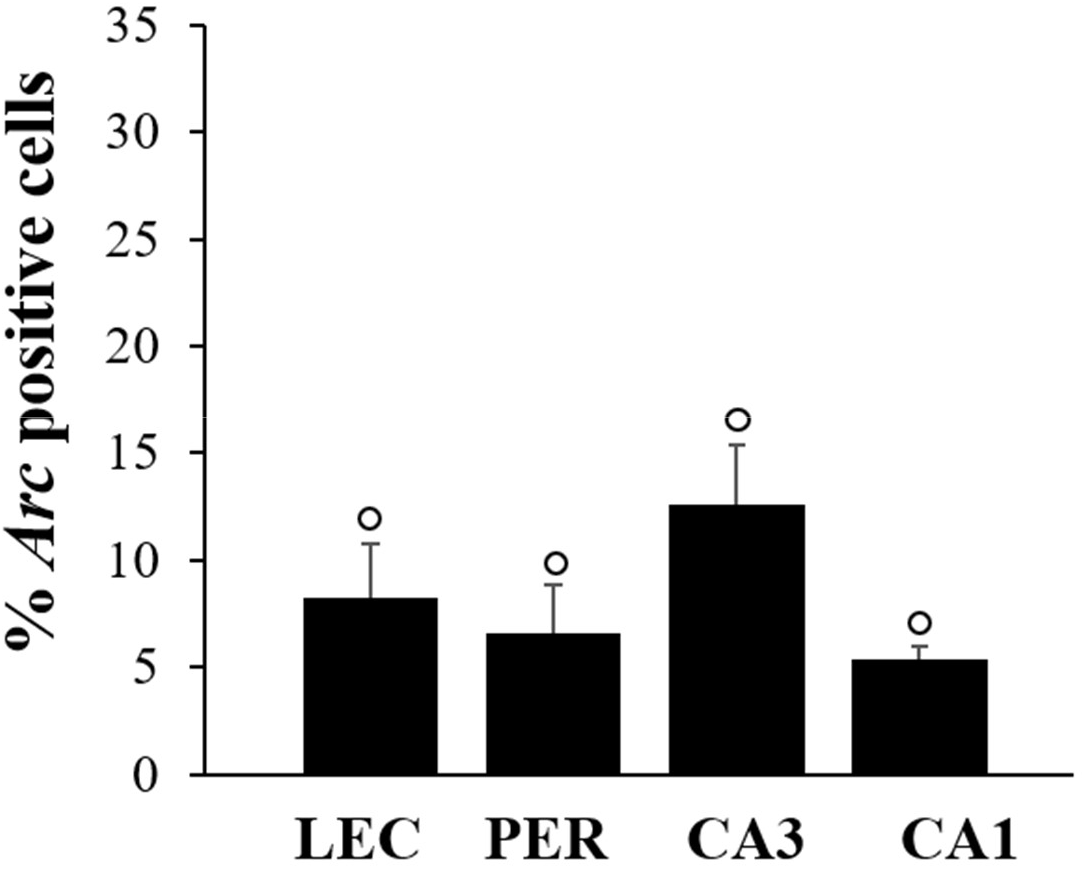
Percentage of *Arc* positive cells in LEC, PER and CA3 and CA1 hippocampal subfields rats that did not perform the task. (means ± SEM). Baseline *Arc* RNA expression in MTL areas was low (comparison to 0: °p < 0.05) and comparable between areas in the home-caged controls.

## Discussion

Here, we have combined a human to rat translational memory task with hippocampal lesions and high resolution imaging that allows for LEC’s activity during recognition memory based on familiarity to be dissociated from PER’ activity. We show that LEC and PER are recruited to a comparable extent when hippocampal function is spared. In addition, we report for the first time that LEC activation is inversely related to hippocampal function, while activation of PER is independent of hippocampal integrity. These results indicate that both LEC and PER contribute to the familiarity process and that their contribution, although quantitatively similar, might rely on different mechanisms.

The fact that LEC is recruited during recognition memory based on familiarity in the present study supports the findings of the only study to date reporting a specific activation of LEC during familiarity judgments, albeit using different experimental conditions^17^. Altogether these results indicate, by extension, that familiarity-induced signals observed in humans in the rhinal cortices do truly involve LEC and not only PER. In addition, we showed for the first time that lesioning the hippocampus enhances selectively LEC’s activity. Importantly though, despite the fact that rats with hippocampal damage performed worse on the DNMS task than rats with intact hippocampus, they were still successful in retrieving memories. This suggests that the increase in LEC recruitment in rats with hippocampal lesion might reflect an attempt at a compensating for hippocampal dysfunction. Such a compensatory mechanism in the form of an increased reliance on familiarity associated with an increased activity in rhinal cortices has been reported in a 3T fMRI study in older subjects with reduced hippocampal function but performance comparable to that of young adults^47^. This latter did however not focus on dissociating LEC’s contribution from that of other part of the rhinal cortex. The technical approach adopted in our study does not enable us to investigate the specific mechanism that leads to the increased activity in LEC. However, it is known that EC and the hippocampus share heavy bidirectional connections, principally excitatory projections but also some long-range GABAergic inhibitory projections believed to play an important role in memory by modulating EC’s interneurons^48, 22, 23, 24^. Hence, it might be reasonable to speculate that the hippocampus could exert a tonic inhibition on LEC in subjects with healthy hippocampus, thereby possibly minimizing the contribution of familiarity to recognition memory and maximizing that of recollection. In turn, damage to the hippocampus might release this inhibition, promoting the use of familiarity judgments and yielding successful memory retrieval, albeit with a reduced performance. Evidence for such a shift in strategy, i.e. from a reliance on recollection to a reliance on familiarity, has been reported in young adult rats following hippocampal lesions and in aged rats in the DNMS task used in the present study^36, 49^. Likewise, patients with MTL damage, including the hippocampus, have been shown to perform better on associative word recognition memory tasks when compound words are used as stimuli i.e. when patients are given the opportunity to unitize the two elements of a pair of unrelated words (example i.e.as in ‘motherboard’) which relies on the familiarity process, instead of retrieving them independently (as in ‘mother’ and ‘board’)^50^. In summary, our results show that LEC plays a crucial role in familiarity and suggest that the HIP to LEC projections might be modulating the extent to which familiarity contributes to recognition memory.

Conversely, damaging the hippocampus did not affect PER activity during familiarity judgments demonstrating that PER activity is not tied to hippocampal function within this frame. The fact that PER and HIP can work independently to detect the familiarity/novelty for items is a well-accepted concept, especially in rodents as shown in lesion and IEG studies using spontaneous object recognition memory task with low memory demands^51 26, 25, 52^. The present study builds upon this existing knowledge and shows evidence that PER and HIP function can also be independent in memory tasks requiring high memory demands (i.e. with a long study list of items, a large delay between study and retrieval phases etc…) and different types of stimuli (i.e. here odors). This underlines the robustness of the findings and the fact that this result is independent of experimental conditions. Also, as mentioned previously, even though the role of PER in familiarity has been extensively studied in humans and rodents^12, 13, 15, 16, 14, 17, 18, 7, 53^, evidence for a specific involvement of LEC has been scarce in animals^17^ and inexistent in humans. Hence, the extent to which the engagement of LEC and PER during judgments based on familiarity is comparable in healthy subjects or when hippocampal function is compromised was not known. Our results address the latter question by showing that LEC and PER’s contribution to familiarity is quantitatively comparable but that they might belong to distinct neuroanatomical subnetworks supporting familiarity judgments as lesions of the hippocampus increased only LEC’s activity within this frame.

Furthermore, as a token of the reproducibility of *Arc* IEG imaging data, LEC and PER’s activity in the group with intact hippocampus were comparable between the present study and the study of Atucha and colleagues^17^, in which the intact hippocampus group was tested under the exact same experimental conditions. Likewise, activity in CA1 and CA3 in the group without hippocampal lesions was also comparable between studies and in line with the findings that rats rely on recollection, supported by the hippocampus^27, 36, 8, 54^, in addition to relying on familiarity to solve the present DNMS task. Notably, in Atucha’s and the current studies, CA3 was more recruited than CA1, possibly reflecting the engagement of CA3 in pattern completion, a mechanism thought to rely on recollection^55^ and taking advantage of CA3’s recurrent collaterals believed to be crucial for autoassociative networks^56, 57^. Also, in the present study, MTL activity levels in rats with or without lesions were found to be tied to the memory demands of the tasks as MTL activity levels of home-cage controls, brought to the experimental room together with the trained animals but remaining undisturbed in their home caged, were very low in comparison. This point was also addressed in the study of Atucha and colleagues, with an additional control group that was exposed to the same experimental conditions as the DNMS rats but was randomly rewarded instead of following a DNMS ruleso that no memory demands would apply. MTL activity levels in the randomly rewarded group were also very low and significantly lower than that of rats with intact hippocampus tested on the DNMS task. This together with the fact that *Arc* RNA expression is closely tied to synaptic plasticity^29, 30, 31, 34^, is reported to better reflect task demands than other IEGs, such as *c-fos* and *zif268*, and not stress levels or motor activity^39, 31, 38, 58^ and is commonly used as a marker of cell activation to map activity in the medial temporal lobe^32, 33, 35^ bring further support to the claim that activity levels in the MTL areas in this study are likely to reflect the memory demands of the task.

Altogether, our results provide clear empirical evidence for the long-standing, yet theoretical, claim of a crucial role of LEC in familiarity. Our findings also reveal that LEC’s contribution is comparable to that of PER, making LEC a main contributor to this process, thus giving further support to the dual process theory predicting that familiarity is supported by LEC and PER^8, 59, 60^. Our data also support to some extent the one process theory^9, 10,11^, according to which the hippocampus contributes to familiarity, but is not rigorously in line with this classical model that predicts that hippocampus might enable familiarity. On the contrary, our results show that LEC’s activity during familiarity judgments is inversely correlated with hippocampal function. Our results complement earlier findings showing that MEC specifically supports recollection in a rat lesion study^61^ and the report that the thinning of EC in elderly affects more familiarity but also recollection judgments in humans^62^, bringing further evidence for a functional dissociation between LEC and MEC in recognition memory. The present experimental approach does not allow for a direct study of the mechanisms underlying LEC and PER’s contribution to familiarity but a well-accepted view in memory research is that representations of distinct items are formed at the level of LEC and PER which, along with back projections to the neocortex, would support familiarity judgments^8^. Our present results allow us to expand on this knowledge and to add that only LEC’s contribution to familiarity is under the modulation of the hippocampus (and not PER’s).

In summary, we report that both LEC and PER function are crucial for familiarity judgments and showed that familiarity signals in LEC are inversely correlated to hippocampal function (but not PER’s) suggesting that brain networks supporting the contribution of LEC and PER to familiarity might be different.

## Acknowledgements

Supported by the DFG Sachbeihilfe SA2146/6-1. The authors would like to thank Jeannette Maiwald for technical help.

## Author contributions

LM, Conception and design, Acquisition of data, Analysis and interpretation of data, Drafting or revising the article; EA, Analysis and interpretation of data, Drafting or revising the article; TK, Drafting or revising the article; MMS, Conception and design, Analysis and interpretation of data, Drafting or revising the article.

